## Supporting Information for "Modulating Nucleic Acid Phase Transitions as a Mechanism of Action for Cell-Penetrating Antimicrobial Peptides"

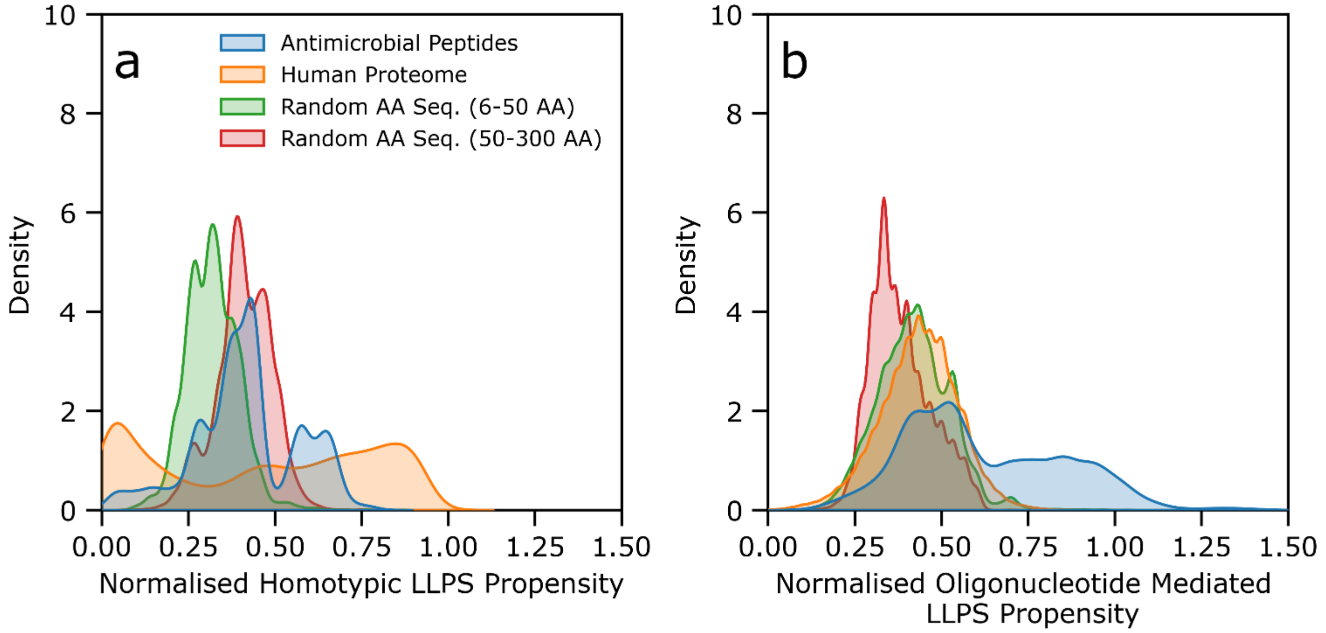


**Figure S1.** Homotypic (**a**) and oligonucleotide-mediated (**b**) phase separation (LLPS) propensity density distribution plots for AMPs (N=13170), human proteins (N=20324), and random amino acid (AA) sequences (N=10000) of different lengths.

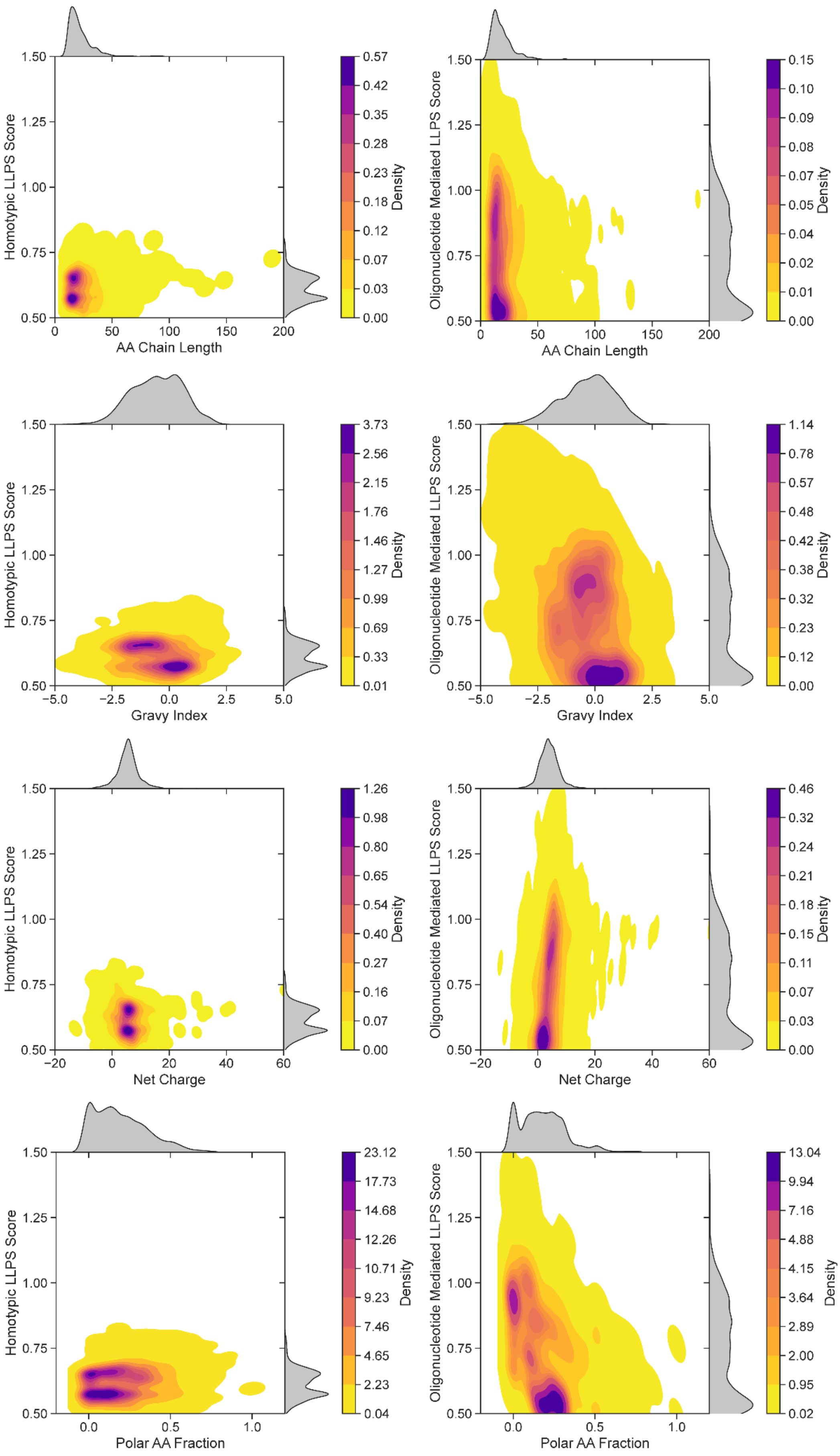


**Figure S2.** Density plots of AMP phase separation (LLPS) score against AMP chain length, net charge, gravy index and polar amino acid fraction.

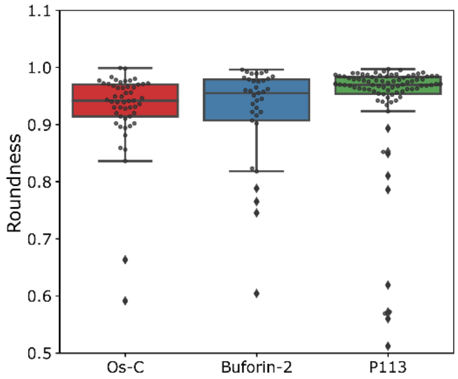


**Figure S3.** Roundness of Os-C-, Buforin-2- and P113-PolyA RNA condensates. Roundness was calculated using the formula: $roundness = \frac{4*\text{area}}{\pi*\text{major\_axis}^2}$. Image analysis was performed using Fiji/ImageJ software. $N_{Os\text{-}C} = 53$, $N_{Buforin\text{-}2} = 38$, $N_{P113} = 89$.

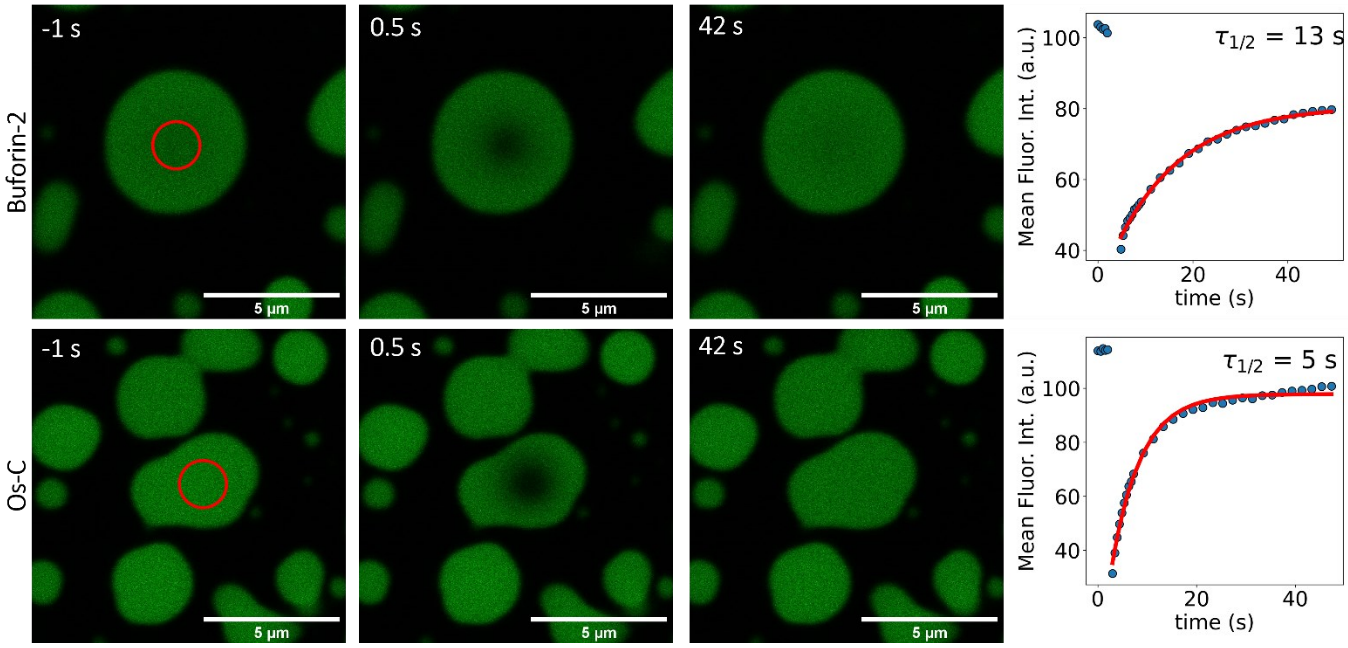


**Figure S4.** Representative confocal microscopy images of FRAP analysis in Buforin-2- or Os-C-PolyA RNA condensates and the kymograph of the FRAP experiment.

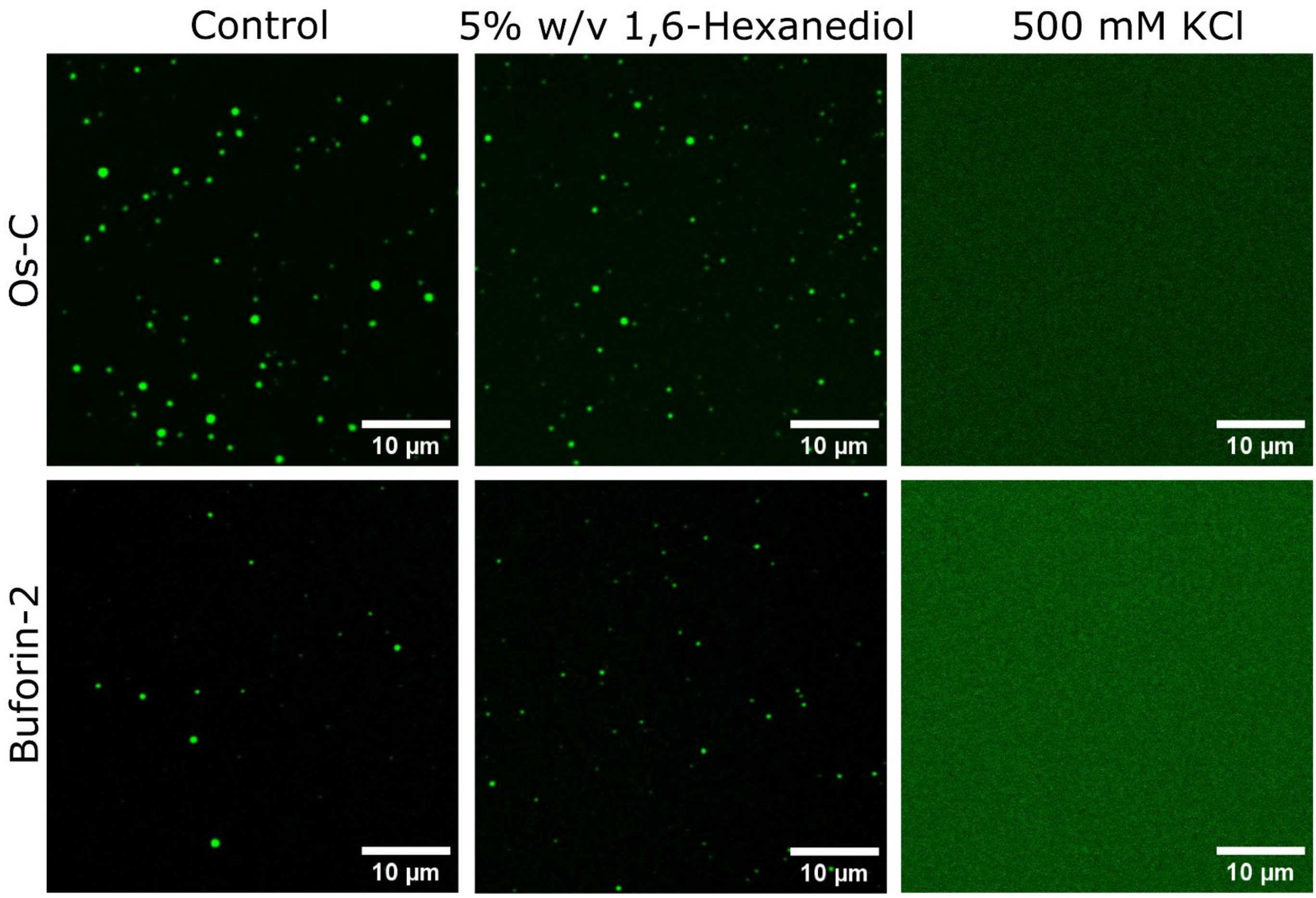


**Figure S5.** Representative confocal fluorescence microscopy images of Os-C- and Buforin-2-Yeast RNA condensates before the introduction of 1,6-hexanediol or a high concentration of KCl.

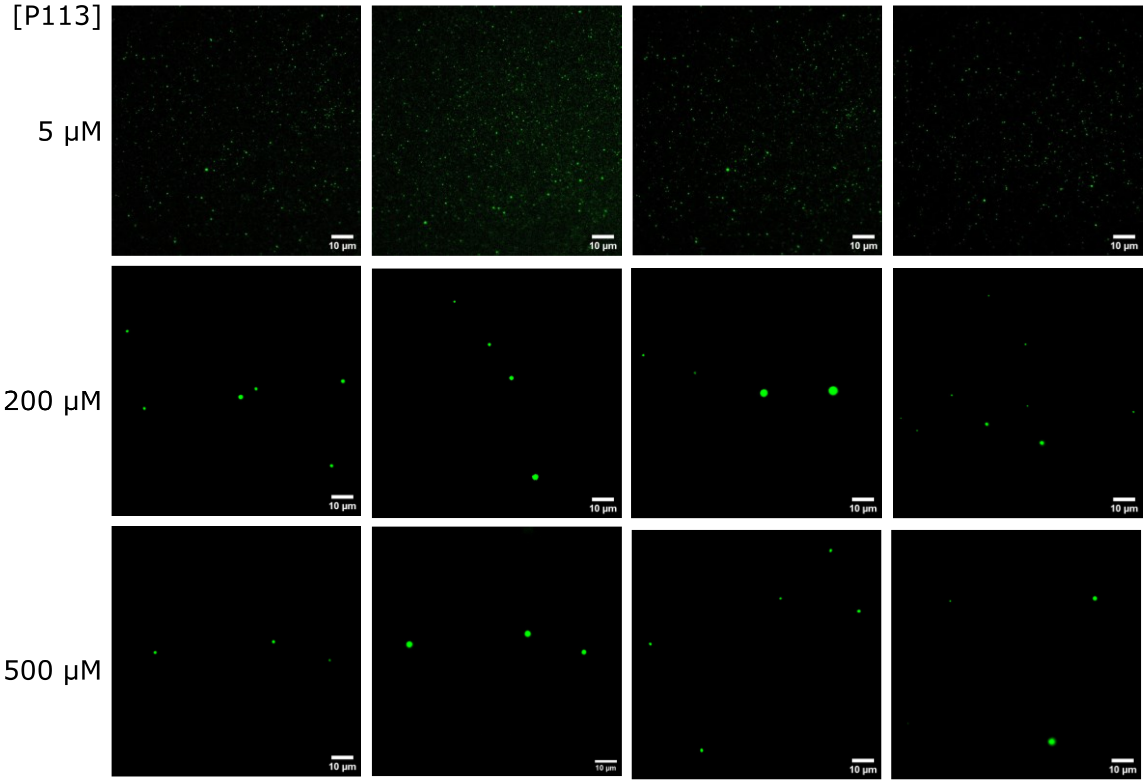


**Figure S6.** Epifluorescence microscopy images of the of Atto 647N-labelled 16S rRNA (0.2 nM final concentration) samples containing varying concentrations of the P113 peptide. Multiple regions of interest are show for every sample.

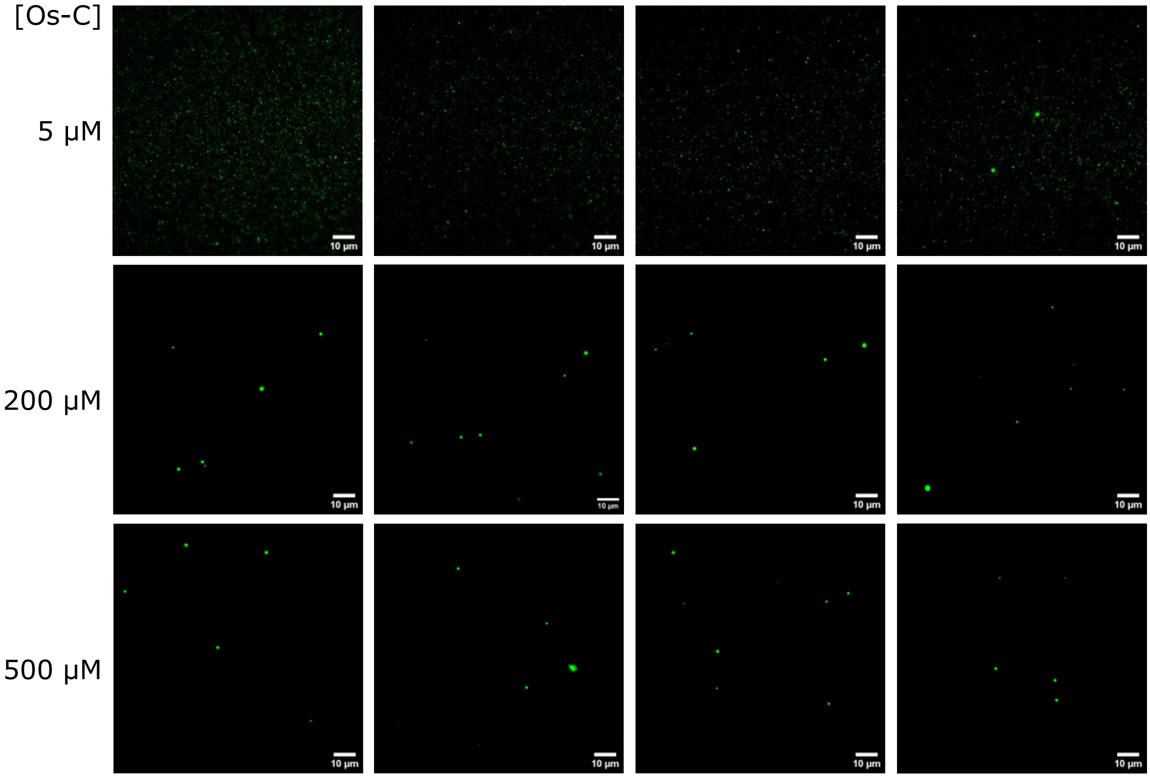


**Figure S7.** Epifluorescence microscopy images of the of Atto 647N-labelled 16S rRNA (0.2 nM final concentration) samples containing varying concentrations of the Os-C peptide. Multiple regions of interest are show for every sample.

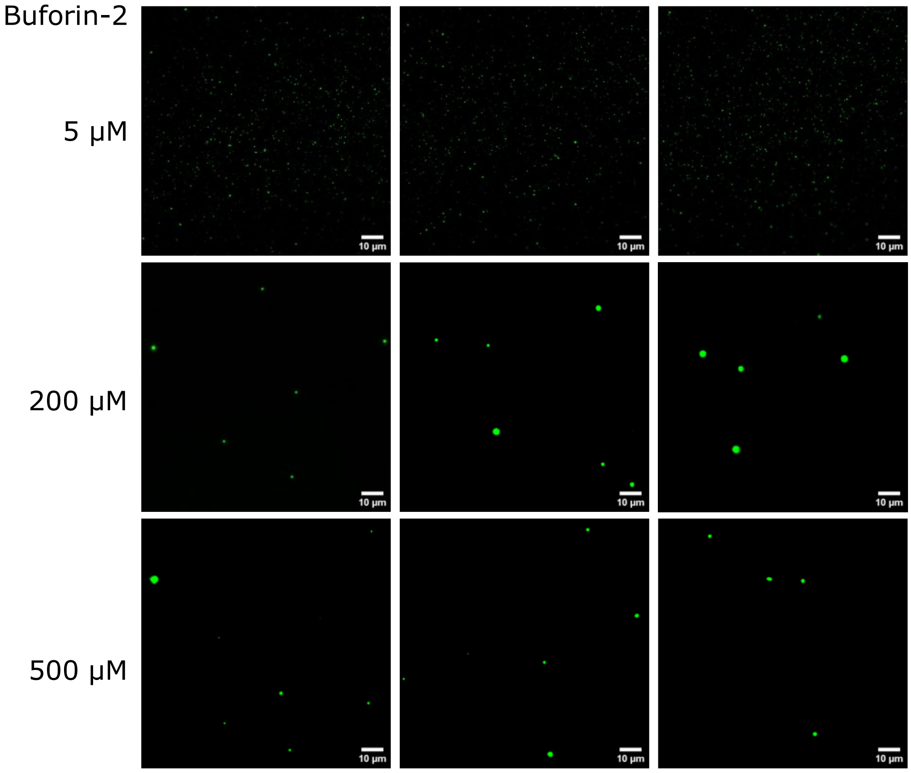


**Figure S8.** Epifluorescence microscopy images of the of Atto 647N-labelled 16S rRNA (0.2 nM final concentration) samples containing varying concentrations of Buforin-2. Multiple regions of interest are show for every sample.

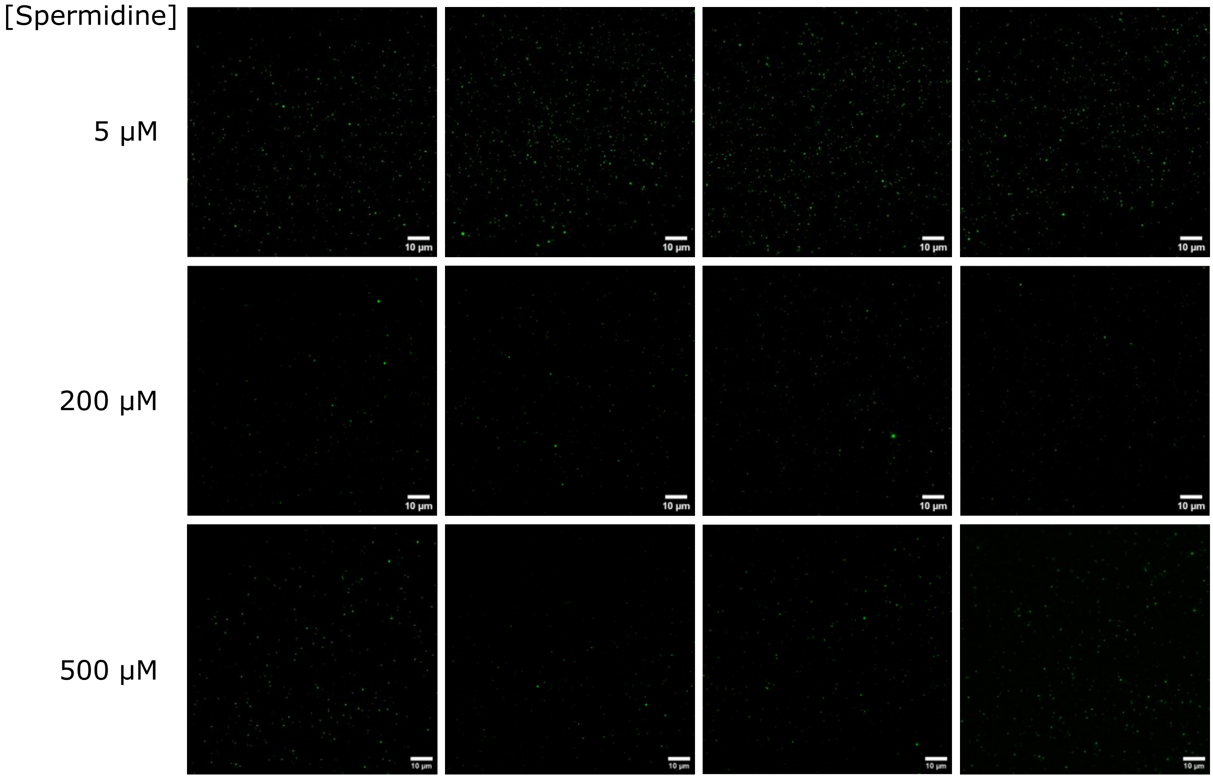


**Figure S9.** Epifluorescence microscopy images of the of Atto 647N-labelled 16S rRNA (0.2 nM final concentration) samples containing varying concentrations of spermidine. Multiple regions of interest are show for every sample.

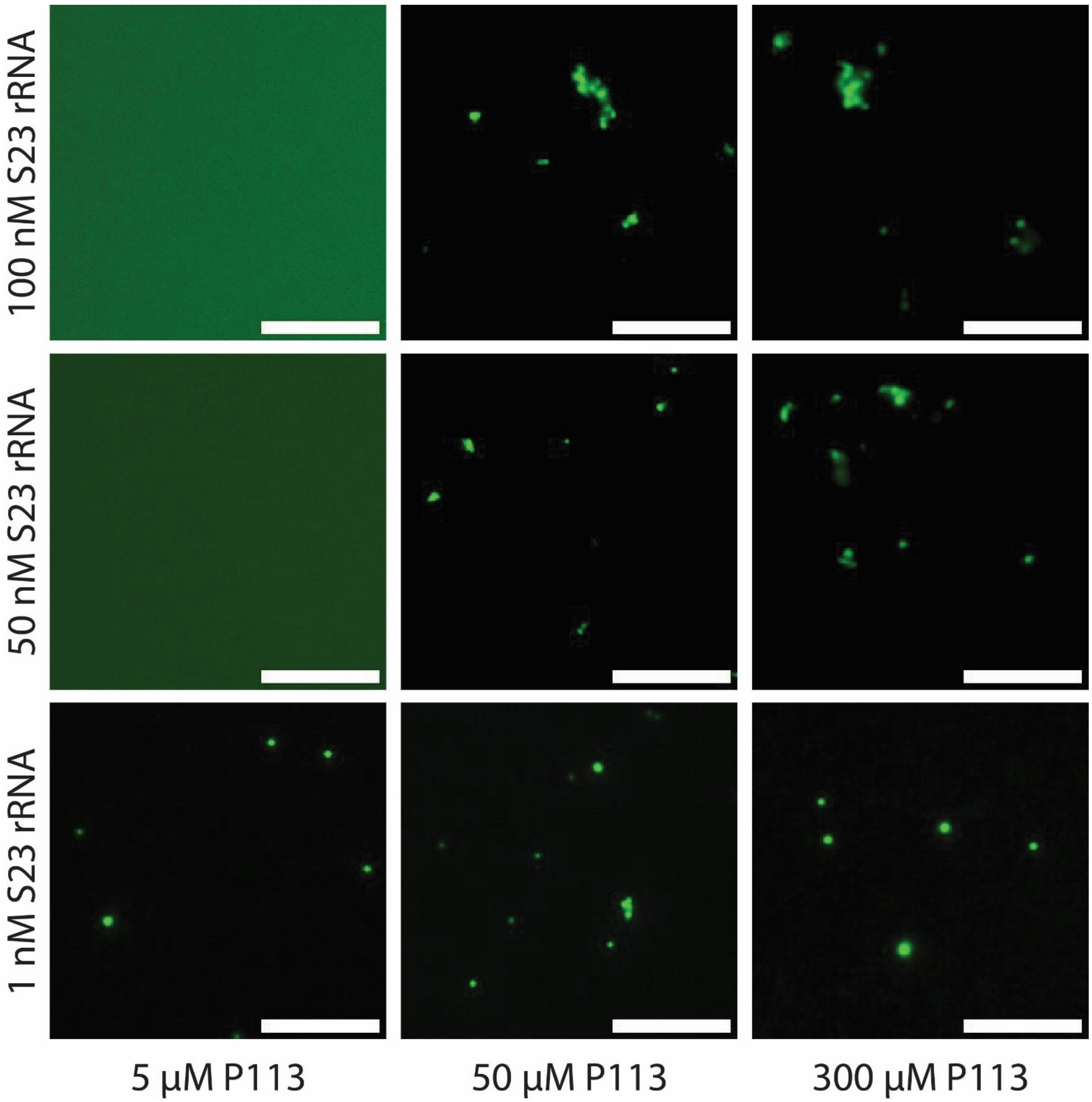


**Figure S10.** Epifluorescence microscopy images of the of Atto 647N-labelled 23S rRNA (1-100 nM final concentration) samples containing varying concentrations of P113.

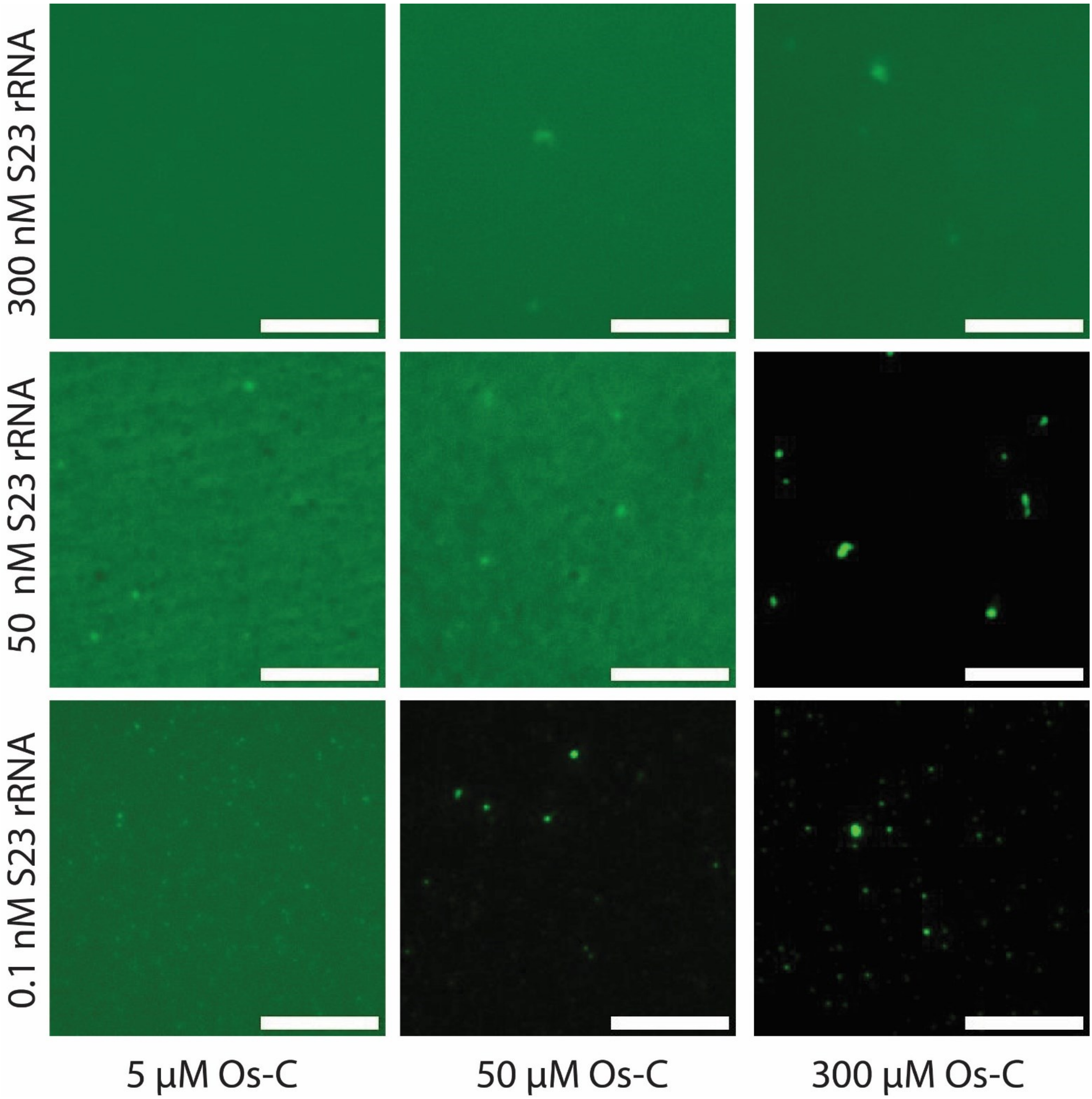


**Figure S11.** Epifluorescence microscopy images of the of Atto 647N-labelled 23S rRNA (0.1-300 nM final concentration) samples containing varying concentrations of Os-C.

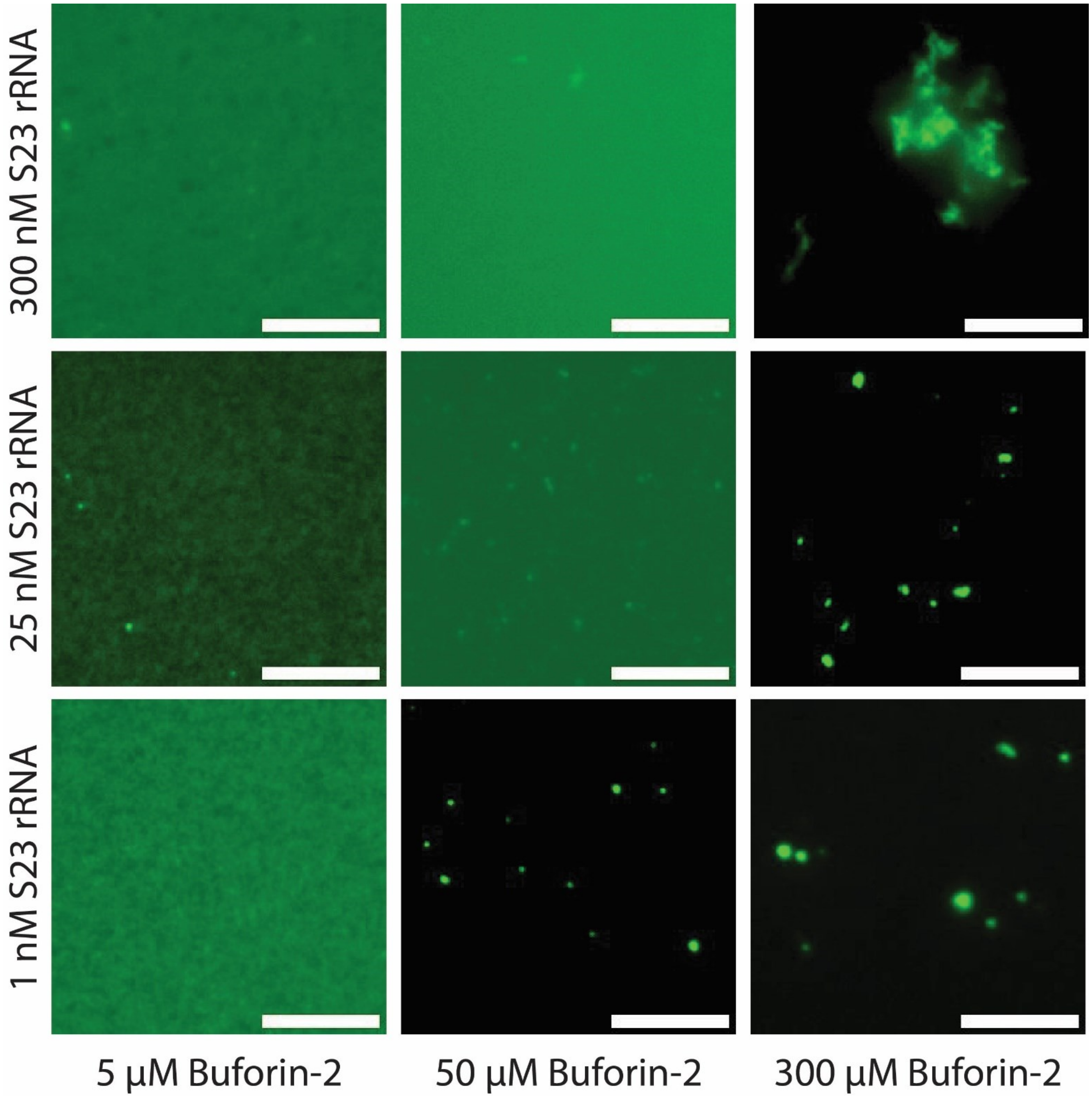


**Figure S12.** Epifluorescence microscopy images of the of Atto 647N-labelled 16S rRNA (1-300 nM final concentration) samples containing varying concentrations of Buforin-2.

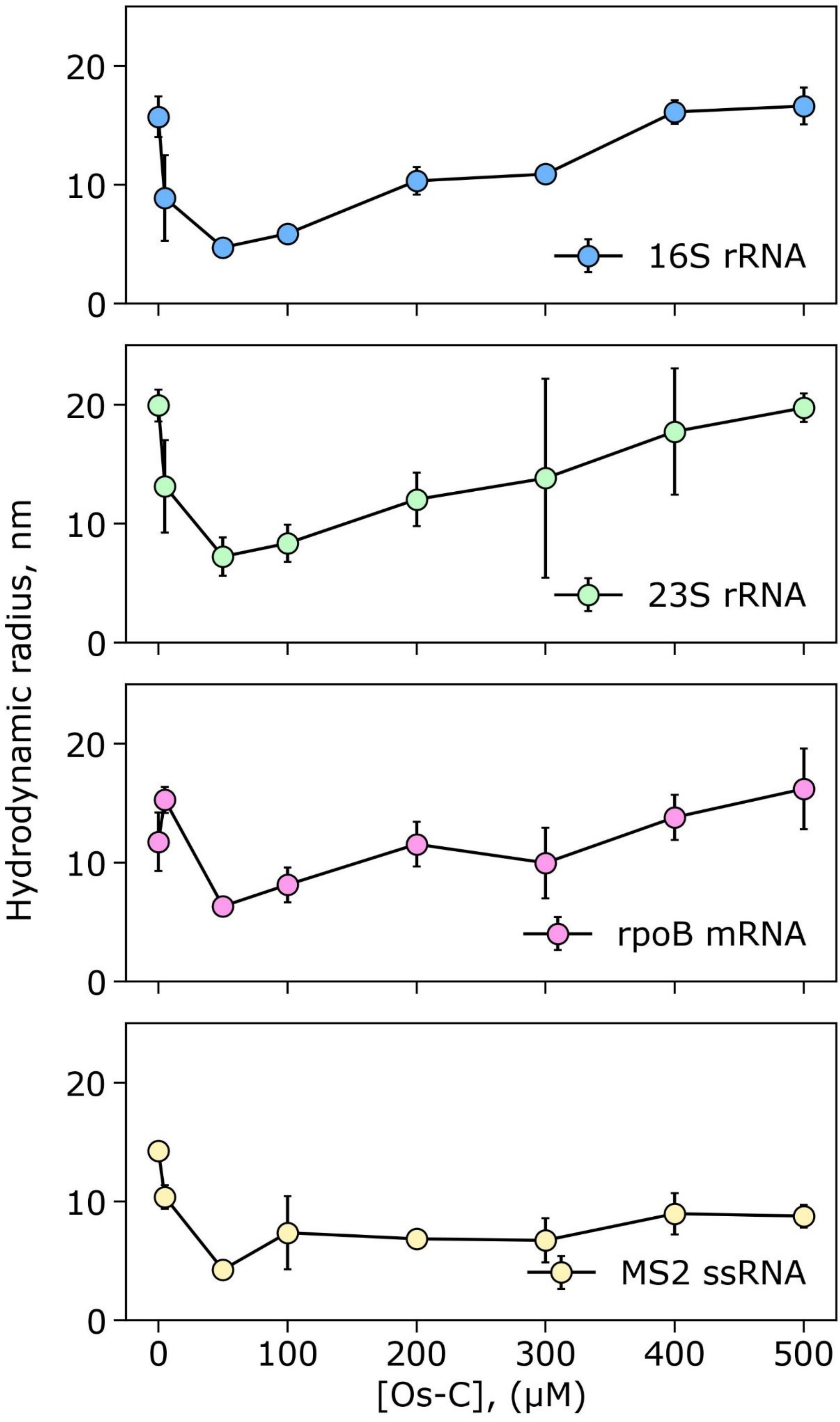


**Figure S13.** MDS measurements of 16S rRNA, 23S rRNA, rpoB mRNA and MS2 ssRNA in the presence of Os-C antimicrobial peptide. Error bars are standard deviations. N = 3.

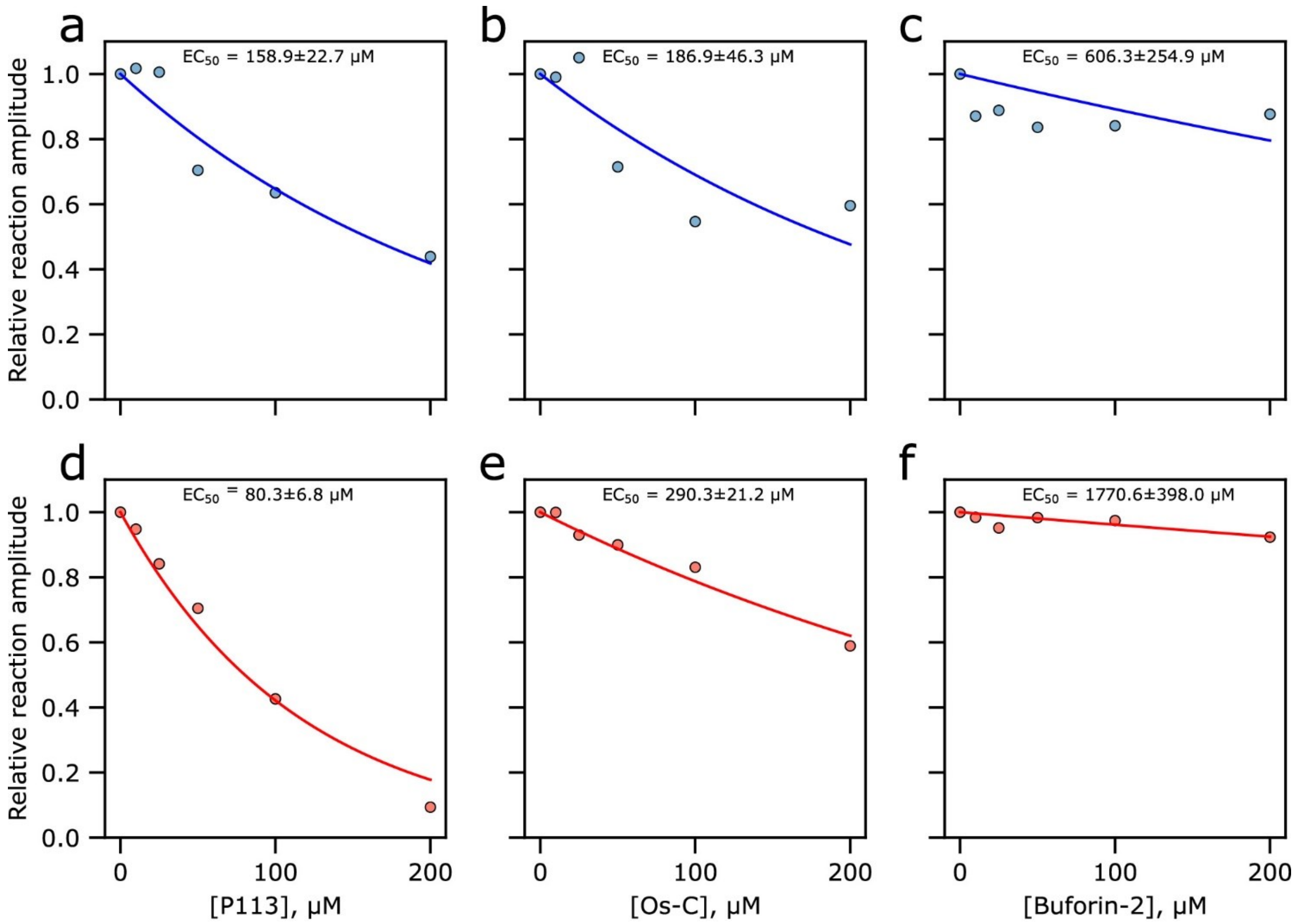


**Figure S14.** Effect of AMPs on relative CFPS reaction aplitude. Relative reaction amplitudes of DNA template-initiated (**a-c**) or mRNA-initiated (**d-f**) reaction. The solid lines are the exponential fits: $y = 2^{\frac{-[AMP]}{EC_{50}}}$, where [AMP] is the concentration of antimicrobial peptide and $EC_{50}$ is half maximal effective concentration. $EC_{50}$ error is the RMS (root mean square).

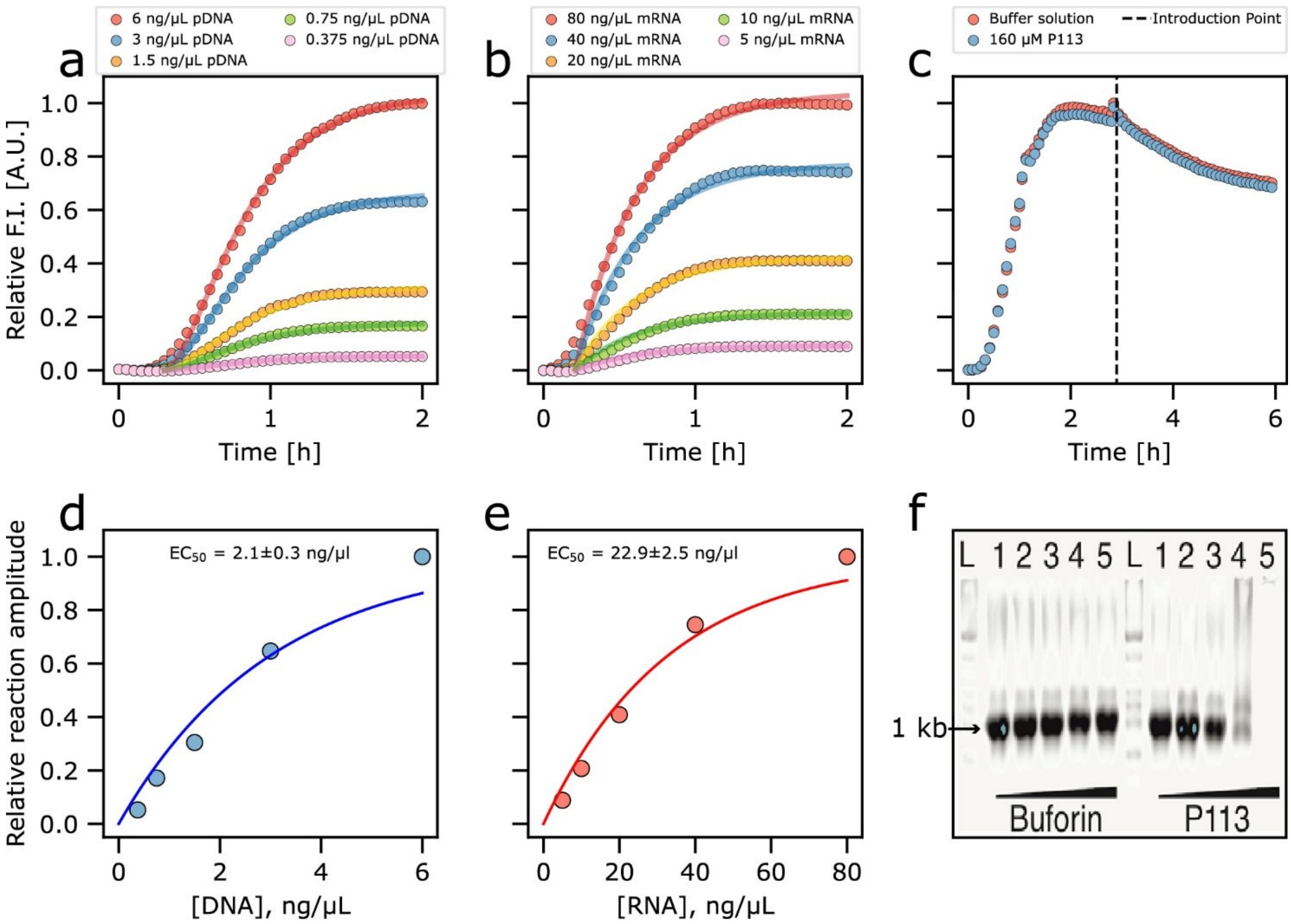


**Figure S15.** Cell-free eGFP expression reaction initiated using plasmid DNA (**a, c**) or mRNA (**b**). Cell-free eGFP synthesis reaction where P113 peptide solution or buffer solution was introduced after 3h since the reaction started (**c**). Relative reaction amplitudes of DNA template-initiated (**d**) or mRNA-initiated (**e**) reaction. The solid lines are the exponential fits: $y = 1 - 2^{\frac{-[NA]}{EC_{50}}}$, where [NA] is the initial concentration of nucleic acid template and $EC_{50}$ is half maximal effective concentration.In vitro transcription reaction in the presence of varying concentrations of Buforin-2 or P113 peptides (f). eGFP transcripts (~1 kb) were produced by adding 25 ng/µl of the template plasmid into T7 polymerase-driven in vitro transcription reaction in the presence of 50-300 µM each peptide. L denotes RNA size marker, lanes 1 – 0 µM peptide (buffer only); lanes 2-5 correspond to 50, 100, 200 and 300 µM peptide concentrations. Note that the additional peptide constituted less than 5% (v/v) of the total reaction volume at the highest concentration shown. P113 significantly inhibits the transcription reaction in the 200 µM range (lane 4).

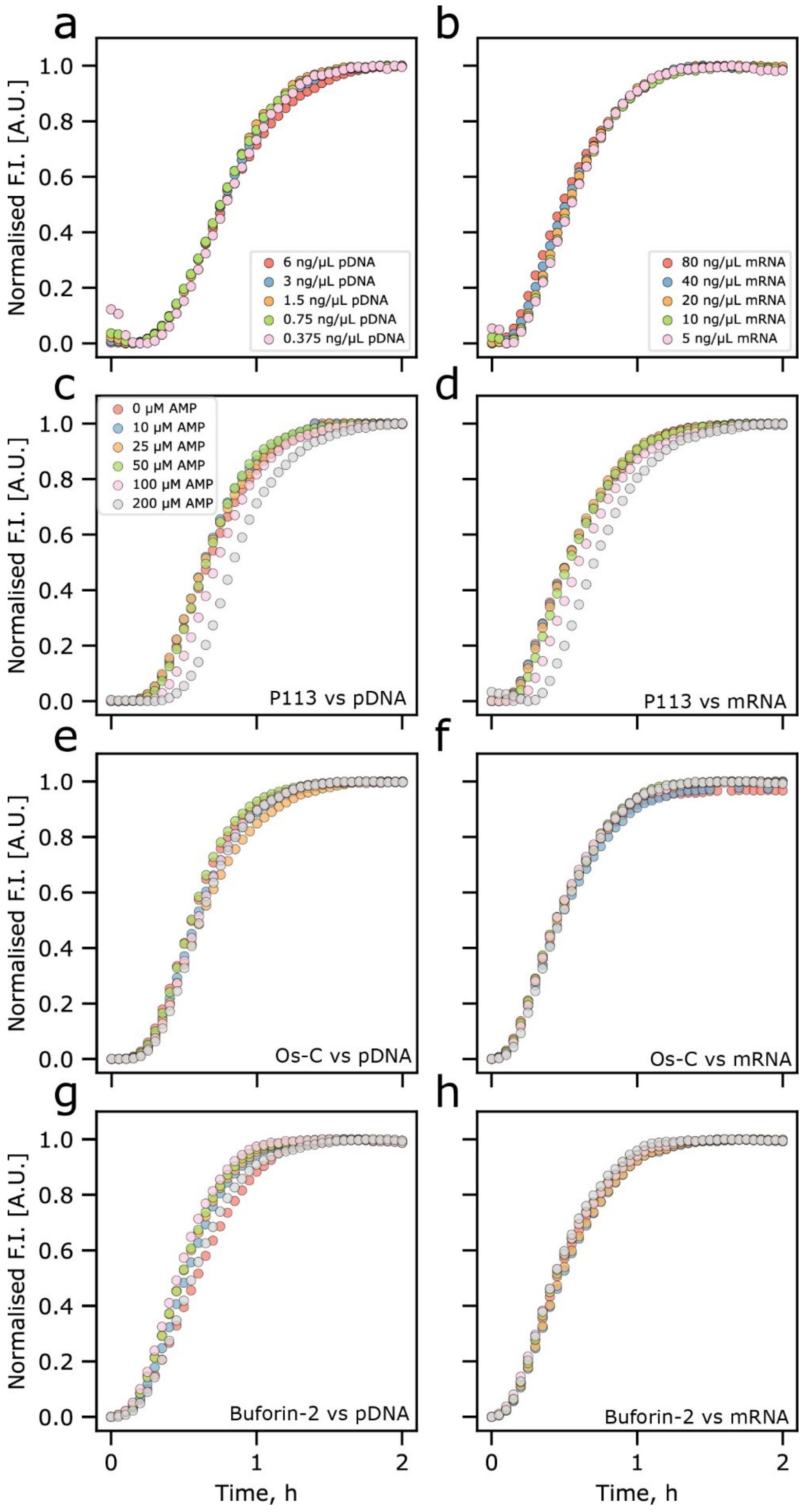


**Figure S16.** Kinetics of cell-free eGFP protein synthesis. Normalised kinetic curves of CFPS reaction initiated by plasmid DNA (pDNA) (**a, c, e, g**) or mRNA (**b, d, f, h**) at varying pDNA (**a**), mRNA (**b**), or AMP (**c, d, e, f, g, h**) concentrations.

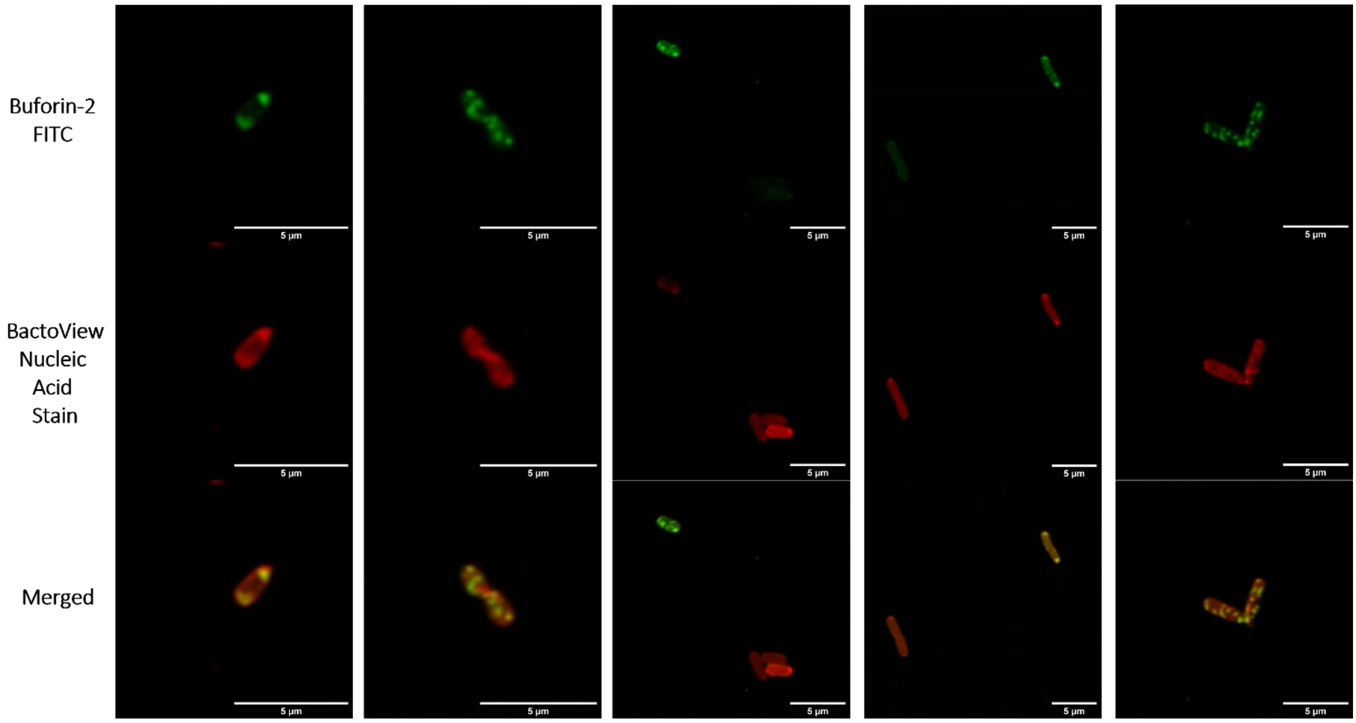


**Figure S17.** Confocal microscopy images of E. coli cells incubated with 20 µM of FITC-Buforin-2 (green) for 2 h. Nucleic acids stained with BactoView Red (red).

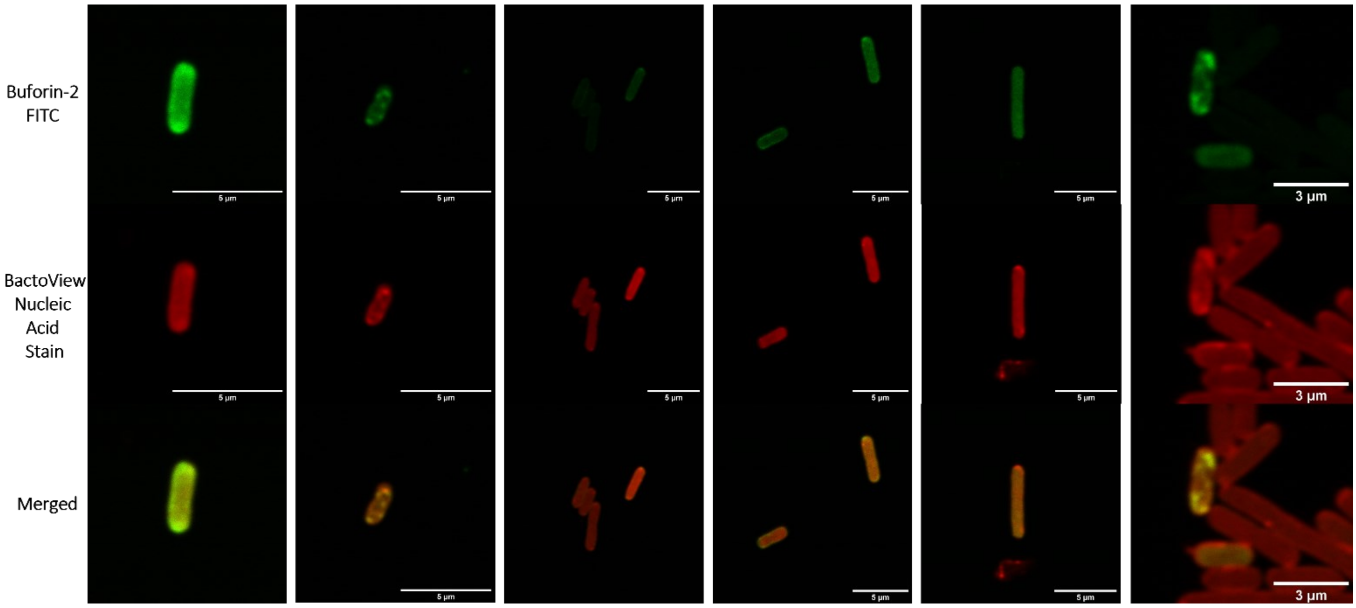


**Figure S18.** Confocal microscopy images of E. coli cells incubated with 20 µM of FITC-Buforin-2 (green) for 2 h. Nucleic acids stained with BactoView Red (red).

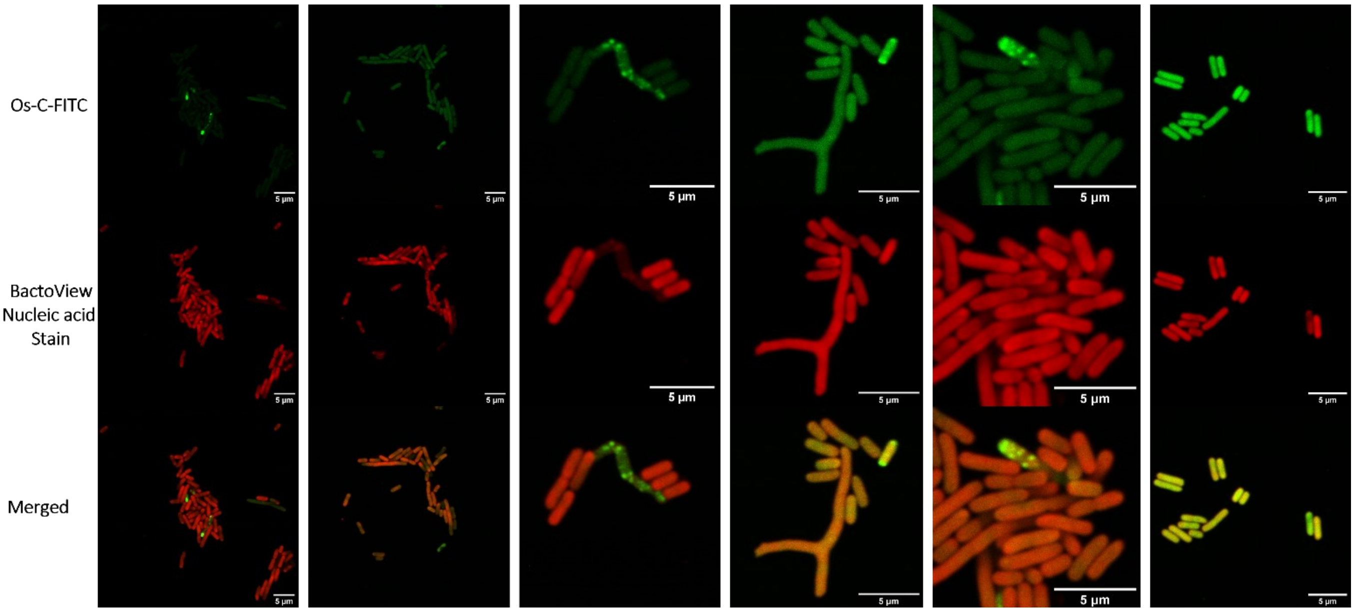


**Figure S19.** Confocal microscopy images of E. coli cells incubated with 20 µM of FITC-Os-C (green) for 2 h. Nucleic acids stained with BactoView Red (red).

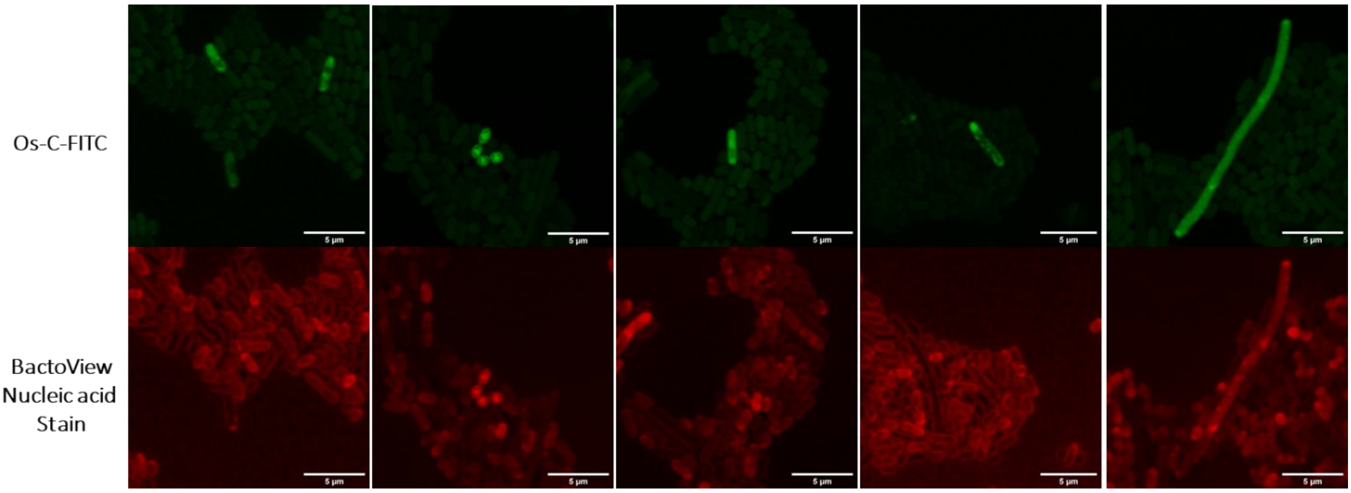


**Figure S20.** Confocal microscopy images of E. coli cells incubated with 20 µM of FITC-Os-C (green) for 2 h. Nucleic acids stained with BactoView Red (red).

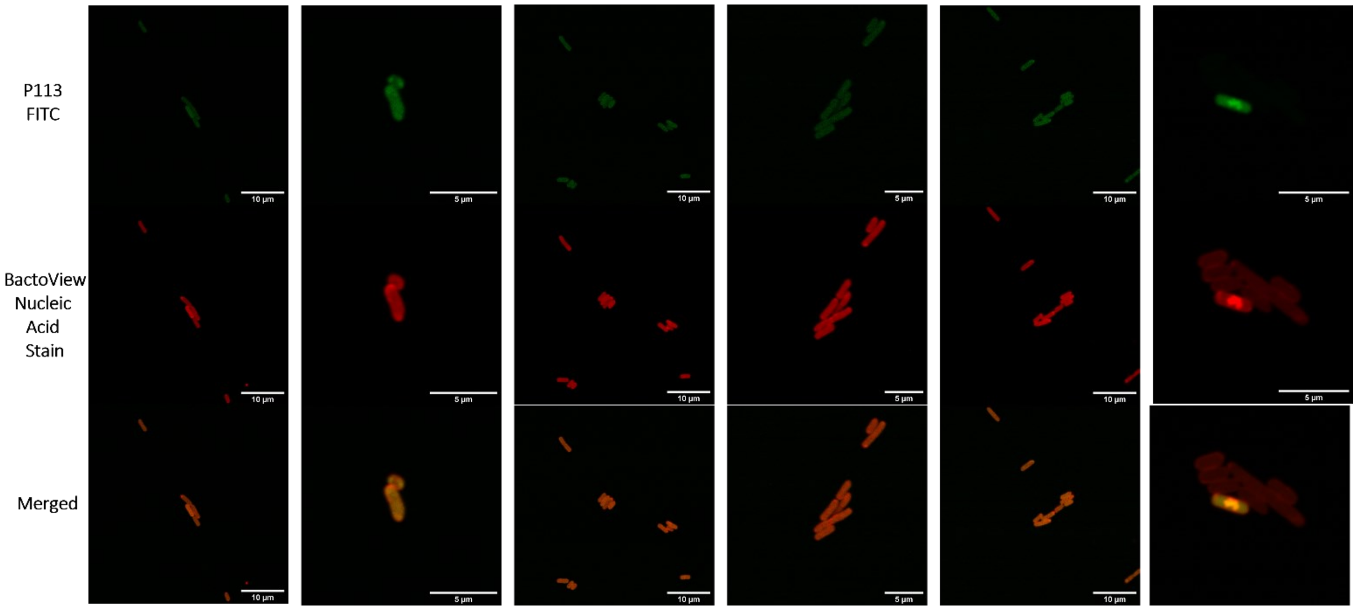

**Figure S21.** Confocal microscopy images of E. coli cells incubated with 20 μM of FITC-P113 (green) for 2 h. Nucleic acids stained with BactoView Red (red).

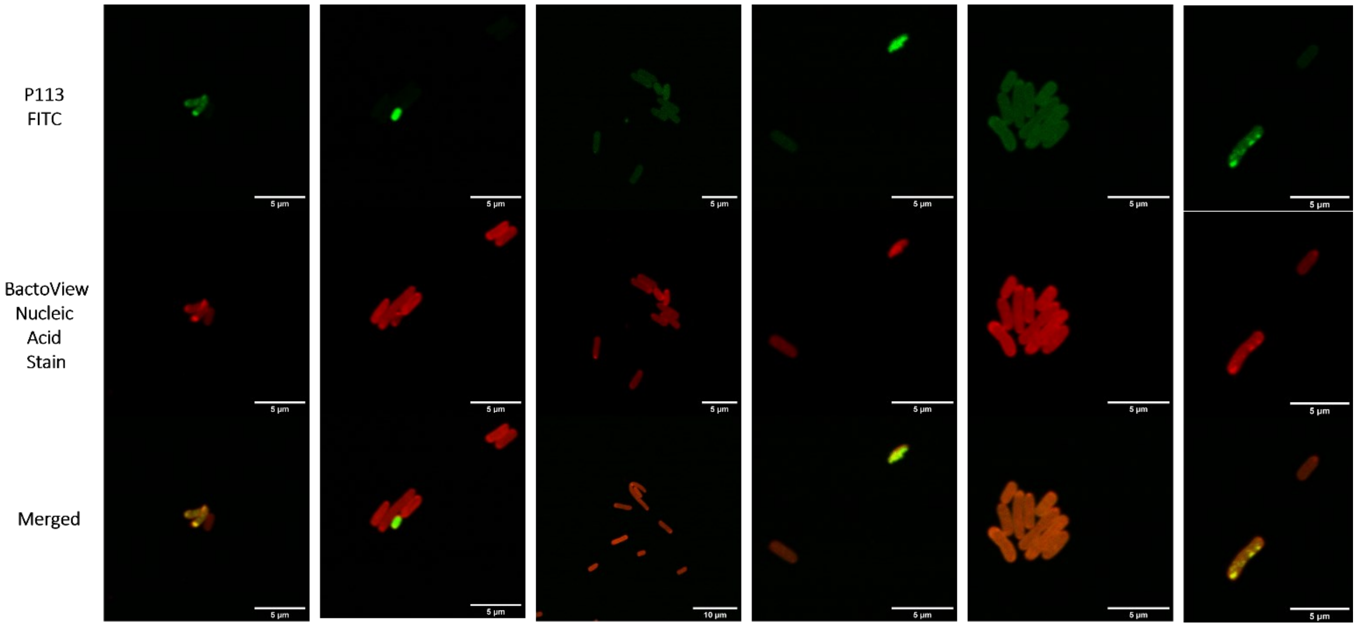


**Figure S22.** Confocal microscopy images of E. coli cells incubated with 20 μM of FITC-P113 (green) for 2 h. Nucleic acids stained with BactoView Red (red).

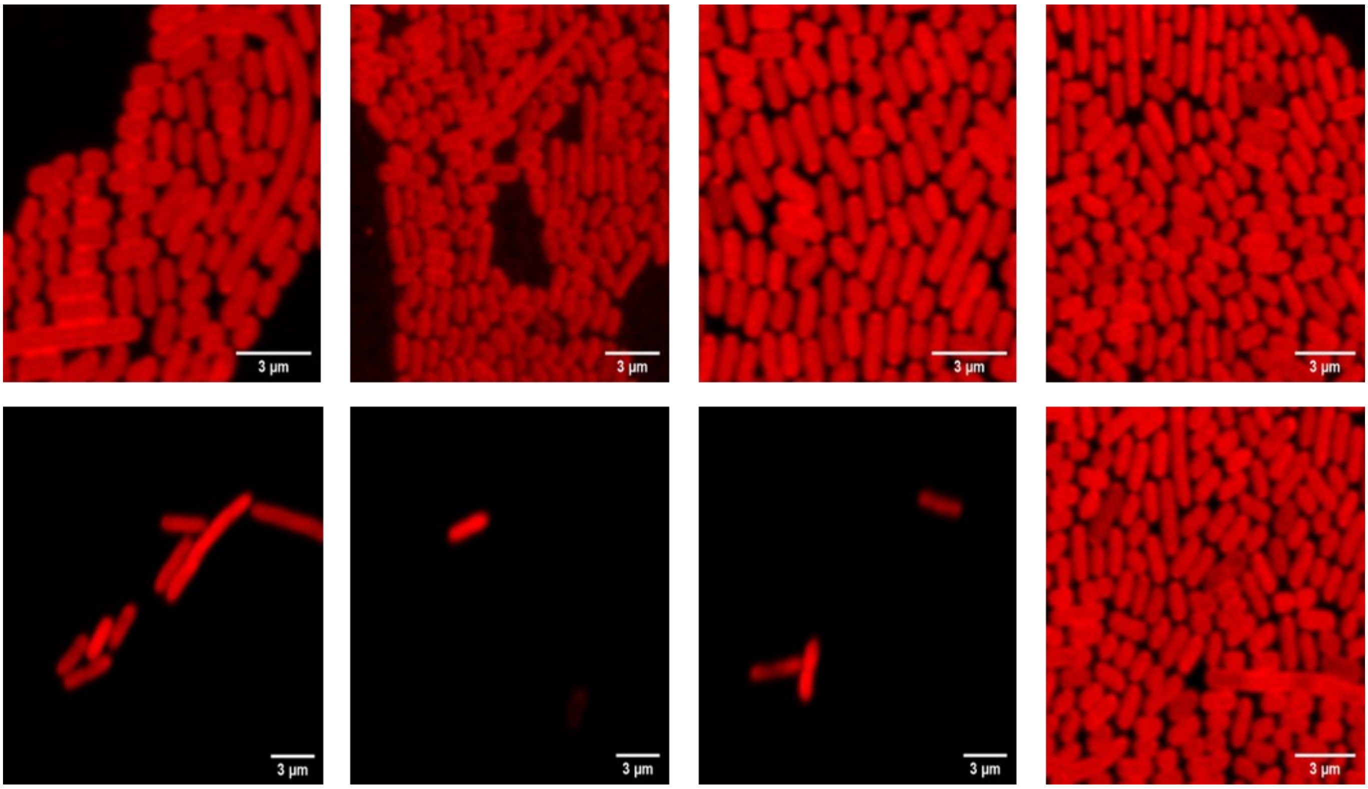


**Figure S23.** Confocal microscopy images of E. coli cells incubated for 2 h without AMP present. Nucleic acids stained with BactoView Red (red). Scale bars are 3 μm.

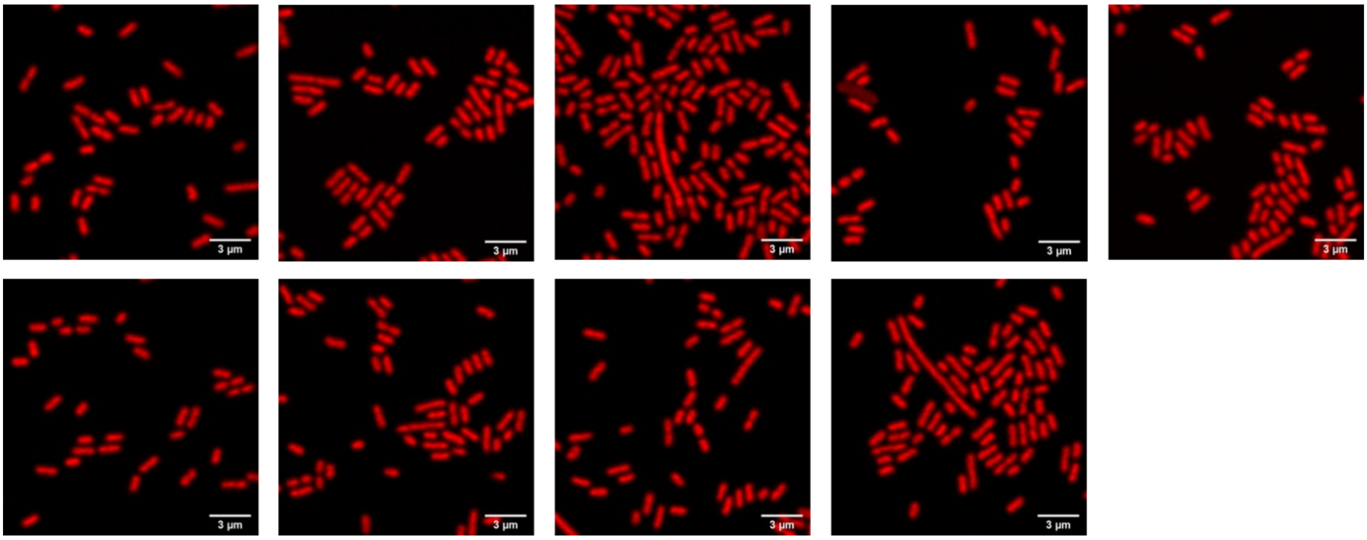


**Figure S24.** Confocal microscopy images of E. coli cells incubated for 2 h without AMP present. Nucleic acids stained with BactoView Red (red). Scale bars are 3 µm.
